# Physics-based Deep Learning for Imaging Neuronal Activity via Two-photon and Light Field Microscopy

**DOI:** 10.1101/2022.10.11.511633

**Authors:** Herman Verinaz-Jadan, Carmel L. Howe, Pingfan Song, Flavie Lesept, Josef Kittler, Amanda J. Foust, Pier Luigi Dragotti

## Abstract

Light Field Microscopy (LFM) is an imaging technique that offers the opportunity to study fast dynamics in biological systems due to its rapid 3D imaging rate. In particular, it is attractive to analyze neuronal activity in the brain. Unlike scanning-based imaging methods, LFM simultaneously encodes the spatial and angular information of light in a single snapshot. However, LFM is limited by a trade-off between spatial and angular resolution and is affected by scattering at deep layers in the brain tissue. In contrast, two-photon (2P) microscopy is a point-scanning 3D imaging technique that achieves higher spatial resolution, deeper tissue penetration, and reduced scattering effects. However, point-scanning acquisition limits the imaging speed in 2P microscopy and cannot be used to simultaneously monitor the activity of a large population of neurons. This work introduces a physics-driven deep neural network to image neuronal activity in scattering volume tissues using LFM. The architecture of the network is obtained by unfolding the ISTA algorithm and is based on the observation that the neurons in the tissue are sparse. The deep-network architecture is also based on a novel imaging system modeling that uses a linear convolutional neural network and fits the physics of the acquisition process. To achieve the high-quality reconstruction of neuronal activity in 3D brain tissues from temporal sequences of light field (LF) images, we train the network in a semi-supervised manner using generative adversarial networks (GANs). We use the TdTomato indicator to obtain static structural information of the tissue with the microscope operating in 2P scanning modality, representing the target reconstruction quality. We also use additional functional data in LF modality with GCaMP indicators to train the network. Our approach is tested under adverse conditions: limited training data, background noise, and scattering samples. We experimentally show that our method performs better than model-based reconstruction strategies and typical artificial neural networks for imaging neuronal activity in mammalian brain tissue, considering reconstruction quality, generalization to functional imaging, and reconstruction speed.

## I. Introduction

**S**TUDYING the rapid dynamics of hundreds of neurons in brain tissue poses a challenge for conventional methods used in microscopy imaging. Typical optical techniques struggle to achieve simultaneous 3D imaging of multiple neurons since they focus on a single plane or point in space. Furthermore, brain tissue is scattering, which increases the difficulty of capturing high-quality images of neurons.

Two-photon (2P) microscopy is an appealing modality for performing 3D imaging in brain tissue. Two-photon microscopy uses near-infrared illumination, which provides deeper tissue penetration and reduced scattering. Furthermore, it restricts excitation to a small volume, mitigating photo-bleaching and providing optical sectioning [2]. As explained in [2], 2P and onwards microscopy has already enabled imaging of neurons in scattering brain tissue. However, due to the limited imaging speed caused by the point-scanning acquisition, 2P microscopy imaging has been largely restricted to planar recordings. On the other hand, light field microscopy (LFM) is a method to image 3D volumes with a 2D camera sensor in a single snapshot. LFM is a fast, non-scanning imaging modality with a volumetric acquisition rate limited only by the camera frame rate, indicator brightness and required signal-to-noise ratio. This performance is achieved by placing a microlens array between the tube lens and the camera sensor of a standard microscope to acquire angular and spatial information from the light simultaneously [3].

In LFM, a 3D image is reconstructed from a 2D light field (LF) image relying on computational reconstruction methods. However, using a single sensor array to capture 3D information limits the reconstruction quality as there is an inherent trade-off between spatial and angular resolution [4], [5]. Thus, recovering a high-quality 3D volume from a single 2D LF image is usually challenging. Furthermore, conventional model-based reconstruction methods for LFM require high computational time. The computation of the forward model and back projection (transpose) usually involves large amount of computation due to the high number of views (hundreds) in the LF data, the lateral upscaling factor of the 3D volume, and the lack of shift-invariance in the system. Even though new model-based approaches have recently emerged to try to address these issues, [6], [7], [5], [8] the computational time required for these approaches still clashes with the primary goal of LFM to reconstruct 3D volume time series.

Reconstruction methods based on deep learning are potential candidates to solve typical problems in LFM [9], [10], [11]. However, current learning approaches are tested under idealized settings that are difficult to achieve in many realistic situations. For instance, when studying neuronal activity in mammalian brain tissue, the sample is highly scattering, nontransparent and contains high background noise, which makes training artificial neural networks (ANN) challenging. On the other hand, model-based optimization approaches have shown to be more robust under these adverse experimental conditions [12], [13].

This work proposes a novel multimodal imaging approach leveraging the respective strengths of 2P microscopy and LFM. We label neurons in the mouse brain tissue using two types of fluorescent proteins: TdTomato and jGCaMP8f. TdTomato captures the static neuron distribution in space, disregarding its activity. On the other hand, the jGCaMP8f is an indicator of calcium concentration which indirectly measures electrical, and therefore functional, activity in the brain. In our setting, we capture the distribution of the neurons in a 400-*μm*-thick brain slice labelled with the TdTomato protein at high resolution using 2P microscopy. Similarly, the LF microscope captures the corresponding LF images for different focal depths. This approach gives us a labelled dataset. In addition, multiple LF temporal sequences are captured from different brain samples using jGCaMP8f protein. The small labelled dataset and a fraction of the LF temporal sequences are used to train our network in a semi-supervised manner, as shown in Figure 1. After training, our network can reconstruct volume time series from LF sequences with high accuracy and speed, despite the temporal LF sequences being obtained with the jGCaMP8f protein, for which we do not have the ground truth volume.

**Fig. 1.**
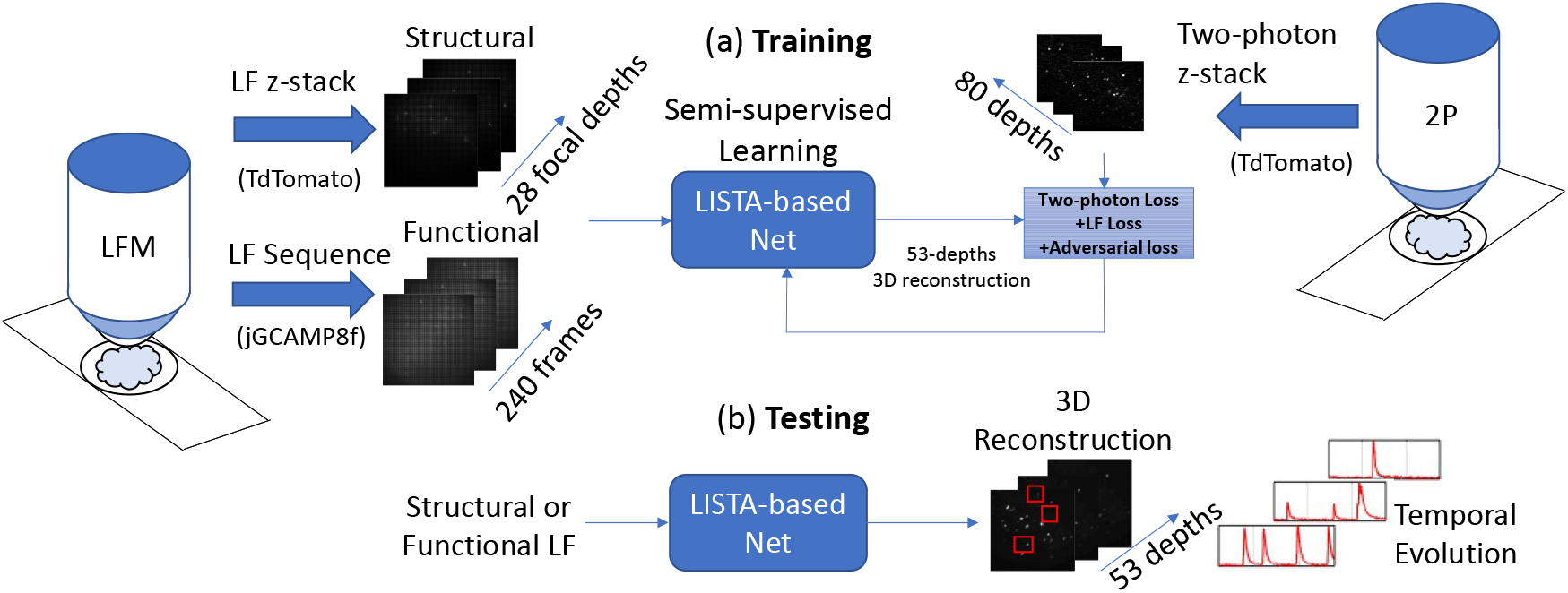
Overview of our approach. As shown in (a), we train our network using a small dataset of 28 training pairs. We effectively collect a 28-slice focal stack of LF images and the corresponding 2P 3D image of 80 depths for a single brain sample labelled with the TdTomato fluorophore. In addition, we use unlabelled data for training. We use three sequences of 80 frames each from 3 different samples for training. These brain samples are labelled with the jGCaMP8f protein which encodes calcium responses in the brain, which is an indirect measurement of electrical brain activity. The architecture of our network is based on the unfolding of the ISTA algorithm. We use a training loss that exploits the knowledge of the forward model and an adversarial regularizer. The testing is performed on LF stacks from unseen samples or LF sequences, and we produce one volume per frame from which we extract neuronal activity, as shown in part (b).

We achieve these results by introducing a physics-driven deep neural network whose architecture is driven by precise modelling of the forward model in LFM and the fact that labelled neurons in tissue are sparse. We leverage the sparsity assumption to design the architecture by unfolding the ISTA algorithm. Finally, a Generative Adversarial Network (GAN) ensures achieving 2P level resolution in the reconstruction while exploiting the knowledge of the forward model, as in model-based approaches. Overall, this approach allows us to exploit the best of two optical techniques to achieve the accurate and fast reconstruction of volume time series of neuronal activity in mammalian brain tissue.

## II. Previous Work

LF imaging has found many interesting applications due to its unique capability of recording the direction of arrival and the location of the light rays. LF imaging has been explored for post-capture refocus, change of point of view, change of focal length, and for robot navigation [14]. Levoy et al. proposed extending LF imaging to microscopy by adapting the Plenoptic 1.0 configuration used for cameras to microscopes [3]. This particular application of LF imaging is interesting because of the capability of capturing volume time series at the camera frame rate, which enables the study of many microscopic biological systems, e.g., networks of neurons in brain tissue.

LFM is a technique that relies on computational approaches to perform the reconstruction of 3D images. Reconstruction techniques in LFM differ from those in photography in that they must consider wave-optics for an accurate system description, whereas the ray-optics model is enough in photography. The first computational approach proposed for reconstruction was presented by Levoy et al. in their pioneering work on LFM [3]. Their algorithm performs digital refocusing to obtain a raw focal stack that is then sharpened by a 3D deconvolution process. Later, Broxton et al. [4] proposed an improvement in the reconstruction by computing the measurement matrix of the system to state a linear inverse problem that is solved using the Richardson-Lucy (RL) algorithm. Currently, large part of the conventional model-based reconstruction strategies rely on similar approaches [15], [16][6][7].

Novel model-based methods have also emerged recently. An artifact-free method faster than conventional RL strategies is proposed in [5]; this approach exploits the low-rank nature of the measurement matrix and uses the Alternating Direction Method of Multipliers (ADMM) to impose additional priors for artifact-free reconstruction. Moreover, alternative model-based approaches exist that only focus on recovering point-like sources, as proposed in [8].

Apart from model-based methods, various approaches that exploit deep learning for reconstruction have been proposed recently. In [9], Wang, et al. describe the first approach that uses an end-to-end convolutional neural network (CNN) for reconstruction. The VCD-Net network is a 2D U-Net trained using synthetic LF data and 3D images obtained with confocal microscopy as labels. The VCD-Net is tested on real LF data by imaging neuron activity in C. elegans and blood flow in the heart of zebrafish larvae. Later, a technique that uses a mixed reconstruction approach was proposed by Li et al. in [17]. The network named deepLFM is designed to enhance the reconstruction obtained after a few RL iterations on LF images. DeepLFM is a 3D U-Net trained and tested using labels obtained by 3D imaging K562 cells with confocal fluorescence microscopy. Then, Page et al. proposed 3D reconstruction from LF images using a network based on a 2D U-net named LFMNet [10]. LFMNet is trained on real LF data and 3D stacks obtained via confocal microscopy. The training data is obtained after imaging brain slices with fluorescently labeled blood vessels. Finally, a convolutional neural network (CNN) named HyLFM that can be retrained to refine the 3D reconstruction with the aid of an additional single plane selective-plane illumination microscopy (SPIM) image has been proposed in [11]. HyLFM is trained on real LF images and SPIM stacks as labels. HyLFM is specifically tested to image medaka heart dynamics and zebrafish neuronal activity.

Works exploring deep-learning methods for LFM show that these techniques are faster and perform better than classic iterative approaches if they are evaluated in controlled scenarios. For instance, they need huge training datasets, low background noise, non-scattering media, or transparent samples. In contrast, model-based approaches are more robust than learning methods and helpful in adverse conditions, as shown in works studying mammalian brain tissue [12], [13]. Thus, we propose fusing appealing features from model-based and learning-based methods. We design an artificial neural network following the physics of the system. The network is trained in a semi-supervised manner by using an adversarial regularizer and exploiting the knowledge of the forward model. Our method achieves high-speed reconstruction after training, and it is robust when imaging scattering samples since it relies on the knowledge of the forward model as in model-based approaches.

## III. Problem Formulation

A LF microscope can be described as a linear operator. In most cases, it is safe to ignore non-linear effects such as occlusion or non-constant refractive indexes of the medium [4], [5]. Therefore, after discretization, a monochromatic (plenoptic 1.0) LF microscope can be represented in matrix form as follows:

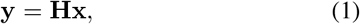

where matrix 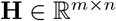 maps a vectorized volumetric input 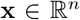 into a LF image 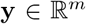. The number *n* of voxels of the volume is usually much larger than the number *m* of pixels of the LF image. In general, the size of **H** depends on the input and output sampling intervals, which are commonly chosen to be *T/s* and *T/N*, respectively (assuming unit lens magnification for simplicity). The constant *T* is the microlens pitch, *s* is an arbitrarily chosen upsampling factor, and *N* is the number of pixels under each microlens (per lateral axis). See Figure 2 for clarification.

**Fig. 2.**
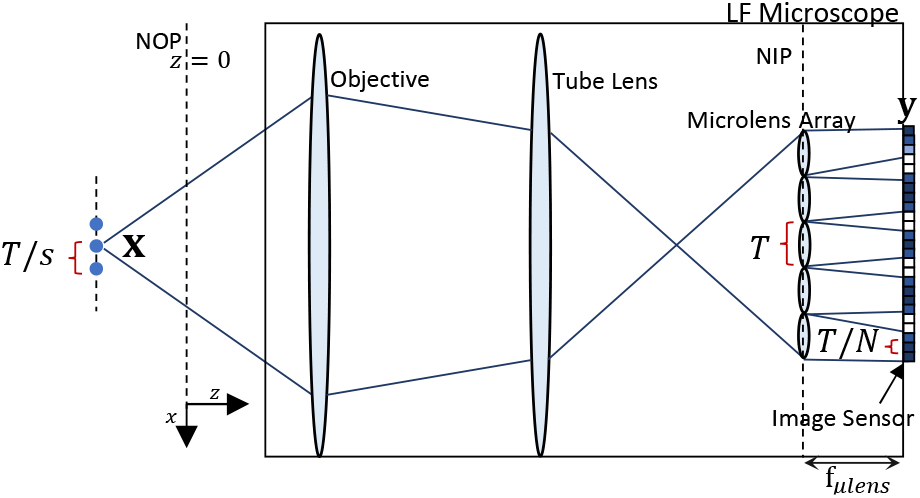
Block diagram of a LF system. In the Plenoptic 1.0 design, a microlens array is placed at the Native Image Plane (NIP) of a standard microscope and the camera sensor is placed at the focal plane of the microlens array. A LF microscope maps a 3D input volume **x** into a 2D LF image **y**. The upsampling factor *s* and the one-dimensional number of pixels under each microlens *N* define the measurement matrix of the system. The optical conjugate plane of the NIP is called the Native Object Plane (NOP).

If one shifts the input of the system laterally by *T*, the output is shifted by *N* pixels, as shown in Figure 3 (a). This behaviour only occurs for shifts which are multiple of *T*. Therefore, for a fixed depth, the forward model is periodically-shift invariant and, therefore, can be modelled by using a filter bank [5], as in Figure 3 (d). Furthermore, the impulse response is unique for each depth, which is the property that allows for localization of sources at different depths, as shown in Figure 3 (b). Thus, to describe the whole system, we need one different filter bank per depth, as shown in Figure 3 (c). The input volume first passes through a slicing operator *S_i_* that selects the depth *z* = *i* for *i* = 1, 2, .., *D*, where *D* is the number of depths. Then, each slice is the input to the corresponding filter bank, i.e., for each *z* = *i*, the *i*-th filter bank outputs a LF image **y**_*i*_. The final output of the microscope **y** is the summation of all the **y**_*i*_. Note that the structure of each filter bank is related to the measurement matrix of the system. Suppose there are *N* × *N* pixels under each microlens and the sampling interval of the volume for both lateral dimensions is *T/s*, in this case, the filter bank structure has *s* × *s* branches, the downsampling factor is *s*, and the upsampling factor is *N*, as shown in Figure 3 (d). As explained in [5], the input and output filter of each branch can be obtained from the measurement matrix **H**. Note that the convolutions and filters are two-dimensional. Similarly, the downsampling factor *s* and upsampling factor *N* refer to both lateral dimensions.

**Fig. 3.**
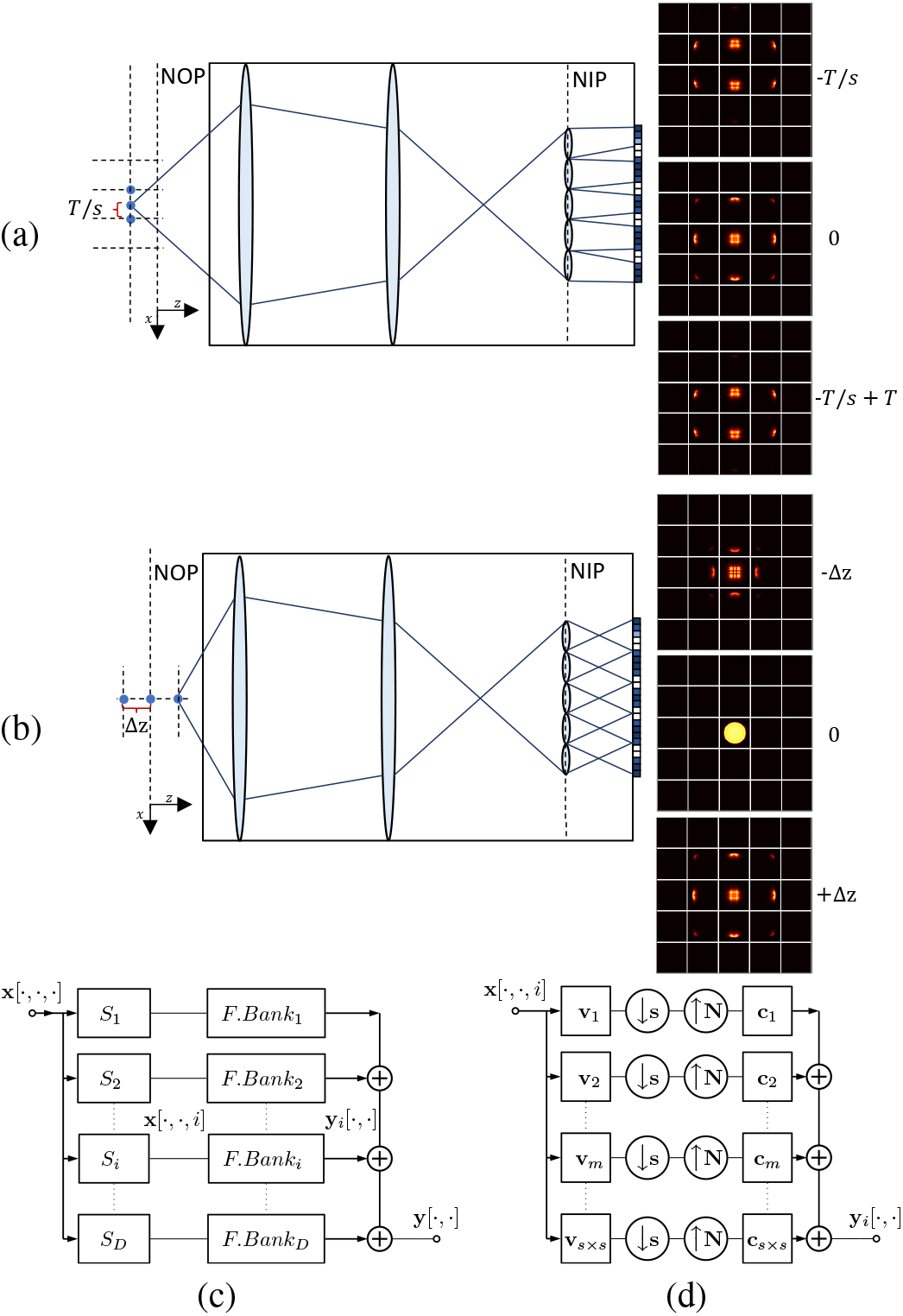
Filter bank representation of the LF forward model. When the input of the system is shifted laterally by a multiple of *T*, the output is also shifted by *T*, as shown in (a). If the input is shifted along the z-axis, the output shows different patterns per depth, as shown in (b). Thus, the output of a LF microscope y can be described as the summation of the output of a group of filter banks. A slicing operator *S_i_* chooses the respective *i*-th depth of the 3D volume, which is the input of the i-th filter bank that outputs a LF image **y**_*i*_, as in (c). As shown in (d), each filter bank has *s* × *s* branches, a downsampling factor of s and an upsampling factor of *N* for both lateral dimensions, where s was chosen arbitrary when computing the measurement matrix **H**, and *N* × *N* is the number of pixels under each microlens.

In this work, we aim to solve the inverse problem derived from Equation (1). Furthermore, studying the spatial and temporal behavior of neurons in brain tissue requires a reconstruction method that performs fast 3D reconstruction from LF sequences. The reconstruction of a 3D volume **x** from a single LF image **y** is traditionally solved using RL-like algorithms. Significant improvements in performance and speed have been achieved with model-based reconstruction, e.g [7], [5]. However, learning-based methods can potentially achieve better reconstruction quality and faster speed when trained properly in controlled scenarios.

## IV. Forward model as a linear CNN

In this section, we propose a novel description of the LF system by using convolutional layers. This description is fundamental for the derivation and implementation of our reconstruction method. We convert the filter-bank model to a linear Convolutional Neural Network (CNN) and propose architectures to perform the forward model computation efficiently.

### A. 4D representation of Light Field

The LF is conventionally represented as a 4D function in photography-related applications. Specifically, the 2D image captured with a LF camera is reordered into a 4D array that is a sampled version of the continuous LF function [18].

Two dimensions of the array represent horizontal and vertical spatial coordinates, while the other two represent horizontal and vertical spatial frequencies. The 4D LF can be interpreted as a collection of sub-aperture images or views, which are 2D images obtained when the spatial frequencies are fixed [18]. See Figure 4 (a) for clarification. In microscopy, the idea of capturing a 4D LF is still valid if the array is interpreted as a sampled version of a 4D Wigner distribution function, a generalization of the concept of LF that considers the effects of diffraction [19].

**Fig. 4.**
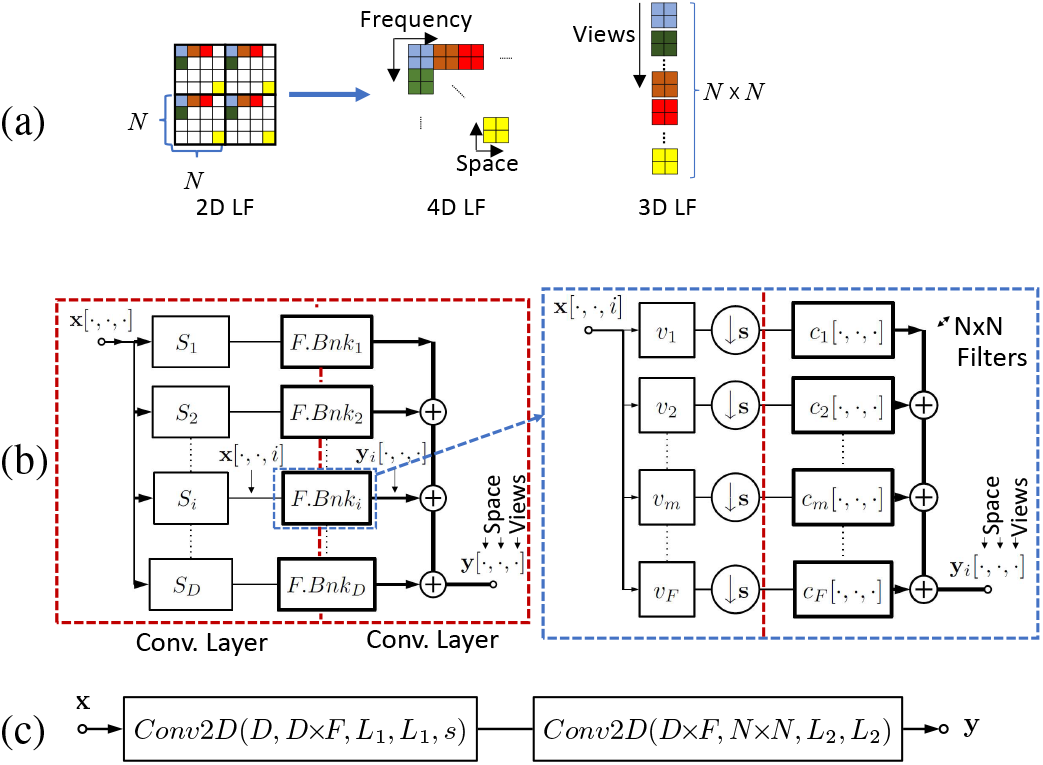
Linear forward model. In (a), we show how a 2D LF can be transformed into a set of *N* × *N* sub-aperture images, where *N* × *N* is the number of pixels under each microlens. In (b), we show how to describe the image formation of a 3D LF image using filter banks. Observe to the right that the model does not contain upsampling blocks since they were absorbed due to the reordering. Furthermore, the synthesis filters were replaced by a group of *N* × *N* filters per branch that introduce the additional dimension representing the view index. In (c), we show the representation of the LFM system as a forward CNN *f*(·). The previous architecture (b) is a particular case of the CNN *f* (·). The notation *Conv*2*D*(·,·, ·, ·, ·) means a 2D convolutional layer with parameters ordered as follows: number of input channels, number of output channels, the height of the filter, width of the filter, and stride. If the stride is omitted, it means unit stride.

### B. Linear CNN

The representation of the LF as a group of views has a convenient property. Unlike the 2D LF image, each view is not an abstract pattern. Instead, it preserves the structure of the original scene since it only carries spatial information. Furthermore, this multi-view representation is attractive for 3D reconstruction in LFM since it is more suitable to fit conventional CNN architectures, as also proposed in [20].

Since the 4D LF can be obtained by just rearranging pixels of the 2D LF, the filterbank description of the LF system described in Section III can be adjusted to explain the formation of a group of sub-aperture images. Specifically, reordering the synthesis filter of each branch allows simple computation of the sub-aperture images, as shown in Figure 4 (b). Note that this new representation follows the basic structure of the original filter bank in Figure 3 (a). However, there is no upsampling block since the original synthesis filter bank is replaced by a group of *N* × *N* filters that leads to multiple outputs forming a set of views or sub-aperture images.

The multiple-view filterbank description of the system can be conveniently implemented using convolutional layers. The LF system can be written as a feed forward network with 2D convolutional layers without bias terms, where the first layer has a stride given by the downsampling factor *s* and the second layer has a unit stride. We represent the CNN by *f*(·) and call it forward CNN (see Figure 4 (c) for clarification). Since the forward CNN is derived from the filter bank representation, the parameters of *f*(·) have a connection with the parameters of the microscope and the physics of the system: the number *N* × *N* of output channels is related to the number of pixels under each microlens, the number of input channels *D* is the number of depths, the filter size *L*_1_ is equal to the downsampling factor *s* or stride [5], the filter size *L*_2_ is instead given by the support of the PSF, specifically, if the support of the PSF related to the largest depth is *M*, then *L*_2_ = *M/N*. Finally, the parameter *F* is related to the upsampling factor. If *F* is set to *s* × *s*, it resembles the theoretical model exactly, while if it is set to a smaller value, it performs an approximation, as explained in [5]. Expressing the forward model as a CNN allows model calibration if a labelled dataset is available. Otherwise, the network parameters can be computed directly from the theoretical model in [4].

### C. Dimensionality reduction

Due to memory and computational-complexity constraints, it is useful to find efficient implementations of the forward CNN. In the previous section, we used two convolutional layers to help map the forward model to the filter banks. However, we can find other alternative architectures that simplify the implementation. First, the 3D input **x** is reshaped to obtain the new input x_*r*_, which has *D* × *s* × *s* channels, as shown in Figure 5 (a). Then, one can replace the two convolutional layers in Figure 4 (c) with a single convolutional layer, as shown in Figure 5 (b). Furthermore, we propose two different simplifications of the architecture based on two observations:

a. Convolutional layers with large filters can usually be well-approximated by a sequence of convolutional layers with smaller filters. Therefore, it is realistic to describe the original system with a series of convolutional layers, as shown in Figure 5 (c). We ensure this architecture uses fewer parameters than the single convolution and also ensure that the size of the equivalent filter is the same as the original one by choosing the filter size *l* and the number of channels *c* accordingly. Note that the weights of the network can be learned from the theoretical matrix **H**. In our work, we use this architecture when we need to apply the forward CNN *f*(·) to a given input volume.
b. The number of channel outputs greatly impacts the number of parameters. For instance, in our setting *N* × *N* = 19 × 19. Therefore, we can reduce the number of views to a smaller value *V* to reduce computational complexity. Then, to restore the original number of views (*N* × *N*), we can add a linear convolutional layer with filters of unit size, as shown in Figure 5 (d). Since the sub-aperture images are usually highly correlated, it is feasible to perform this linear approximation without significantly impairing the accuracy of the model. Note that the input convolutional layer in Figure 5 (d) is named *h*(·). This layer is named compressed forward CNN and is connected to our reconstruction approach.

**Fig. 5.**
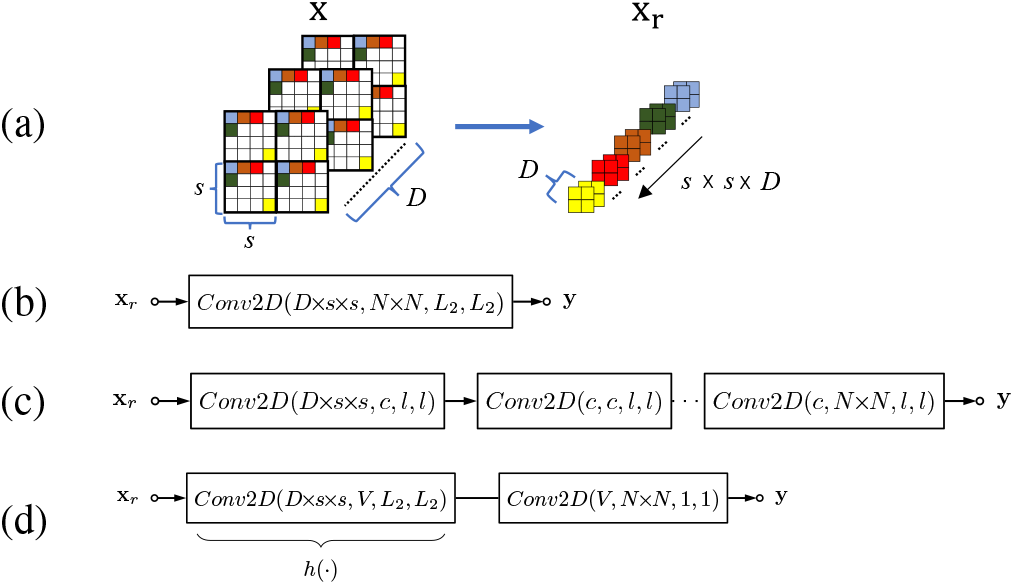
Simplified forward model CNN. We show three different linear CNNs architectures that transform a 3D input image x with *D* depths into a LF image *y* with *N* × *N* views, where *N* × *N* is the number of pixels under each microlens. In all the cases, the input **x** is first reshaped, as shown in (a). The model in Figure 4 (c) can always be converted into the single convolutional layer shown in (b). In (c), we approximate the single convolutional layer by a sequence of convolutional layers with filters of smaller size *l*. Finally, a second approximation is shown in (d), where the first layer *h*(·), named compressed forward CNN, outputs *V* channels, and the second layer recovers the *N* × *N* views with filters of unit size. The notation used for the convolutional layers is the same as in Figure 4(c).

In the sequel, we use *f*(·) to compute the forward model, while the compressed forward CNN *h*(·) is used in the reconstruction network. This will be clarified in the following section.

## V. 3D Reconstruction

In this section, we design a CNN that considers the physics of the system to perform the reconstruction. The architecture of our network is constructed using the unfolding technique [21] and is obtained by unrolling sparsity-driven algorithms for reconstruction. Furthermore, our network is trained in a semi-supervised manner to alleviate the lack of data which is a typical issue for applications in neuroscience.

### A. CNN architecture

Large distributions of labelled neurons can be modelled as compact cell bodies sparsely distributed in brain tissue [22]. Therefore, to reconstruct high-quality 3D volumes, we can consider the following optimization approach that promotes sparsity in the reconstruction:

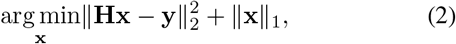

where **y** is a given LF image, **x** is the reconstructed volume. This problem can be solved using the Iterative Shrinkage-Thresholding Algorithm (ISTA) [23] by computing at each iteration:

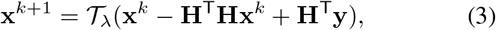

where 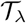 is the soft-thresholding operator with parameter λ. One can interpret each iteration of ISTA as a layer of a neural network with fixed weights. Therefore, it is possible to design a neural network architecture based on ISTA. LISTA [21] (the learned version of ISTA) is a neural network built such that each layer corresponds to one iteration of ISTA. Effectively, each layer of LISTA implements the following step:

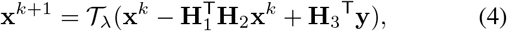

where **H**_1_, **H**_2_ and **H**_3_ are matrices of same size and structure as **H**. These matrices are the parameters of the network that can be learned using a proper loss function. Note that, contrary to [21], we do not fuse the product 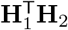 into a single matrix since we want to keep the structure of each factor. This version of LISTA uses the soft-thresholding as the element-wise non-linearity due to the *l*_1_ constraint in Equation (2). However, ISTA can be used with different types of non-linearities related to the prior imposed, as explained in [24]. For instance, replacing 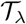 by a rectified linear unit (Relu) imposes non-negativity, and replacing it with a ReLU with a bias term imposes sparsity and non-negativity. In our case, **x** is sparse and non-negative. Therefore, we propose a LISTA network that uses a ReLU with a bias term as non-linearity:

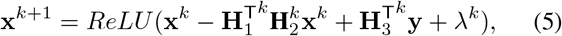

where λ^*k*^ is a learnable bias. Furthermore, the custom 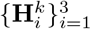 for each unfolded iteration *k* gives the network more capabilities without compromising its simplicity.

In many practical cases, the described LISTA network cannot be used directly to solve the volume reconstruction problem. The size and structure of the matrix **H** make it computationally prohibitive to perform matrix multiplications repeatedly. Therefore, we propose using the compressed forward CNN *h*(·) proposed in Section IV-B to reduce the computational complexity. The final architecture of our network is, therefore, described as follows:

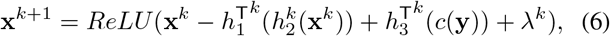

where we have replaced matrices 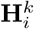 in Equation (5) with the linear mappings 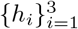. The computation of all the 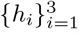 is determined by the architecture of the compressed forward CNN derived from physics and explained in Section IV-B. Note that the structure of the adjoint operators (transpose) 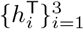 in Equation (6) can be easily computed from the permutation of the weights of *h*(·). Furthermore, the input of the network is *c*(**y**) rather than **y**. The mapping *c*(·) is defined as a single linear convolutional layer with *N* × *N* input channels and *V* output channels and filters of unit size. By having *V* output channels, *c*(·) is compatible with the input size of the operators 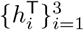. For this compression step, we found unit-size filters to be effective; however, filters of any size could be used. We highlight that the coefficients of the compression layer *c*(·) are learned together with LISTA. The end-to-end network *g*(·; *θ*), where *θ* represents the learnable parameters of the network, is shown in Figure 6. If additional simplification is needed, some convolutional layers in *g*(·) can be replaced by series of convolutional layers with smaller filters.

**Fig. 6.**
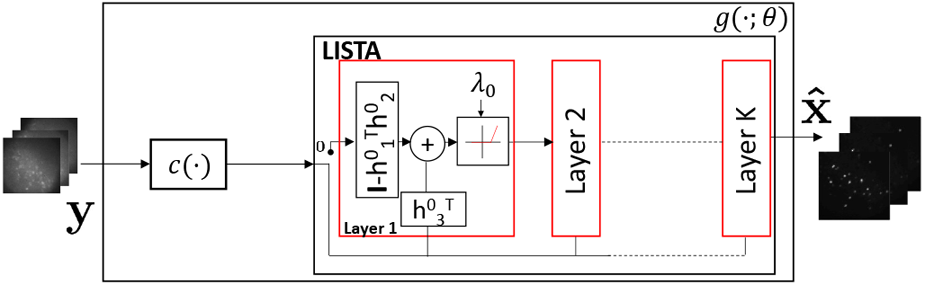
CNN architecture. Our reconstruction network *g*(·) is composed of (1) a compression layer *c*(·), which is a linear convolutional layer with *N* × *N* input channels and *V* output channels and (2) a LISTA network. At each layer of LISTA we use the architecture of the compress forward CNN *h*(·) shown in Figure 5 and the adjoint operator *h^T^*(·). The LISTA network is composed of *K* layers.

### B. CNN Training

We learn the parameters θ of our LISTA network *g*(·; *θ*) with a proper loss function and a mixture of labelled and unlabelled datasets. In our scenario, a labelled dataset comprises LF images and the corresponding 2P volumes. For many applications in LFM, capturing a huge labelled dataset is too expensive or even unfeasible. For instance, when studying the behavior of neurons in mammalian tissue, capturing a clean 3D label is challenging due to the scattering media. Furthermore, using only synthetic data for training is problematic if noise is not appropriately modelled.

In our setting, we propose acquiring a very small labelled training dataset. We label neurons in a single brain sample using TdTomato fluorophore. The TdTomato allows capturing the static distribution of the neurons in space using both 2P and LF modalities. The 2P raster scanning modality provides the ground truth volume that can be paired with the LF images acquired with the same fluorophore. Therefore, to train LISTA we exploit the small labelled dataset, the large amount of unpaired LF images, and the knowledge of the forward model. The training loss is stated as follows:

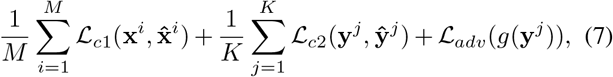

where **x**^*i*^ is the 2P 3D image, 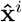 is the network reconstruction, **y**^*i*^ is a LF image, *M* is the number of 3D samples, *K* is the number of LF samples and 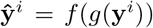, where *f*(·) is the known forward CNN. Notice that the operator f (·) is fixed since it is already known and is based on the model [4]. The loss 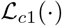 is the 2P content loss computed on the labelled dataset. The loss 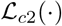 is the LF content loss, which ensures that the re-synthesized LF computed from the recovered volume is close to the original LF image. The adversarial loss 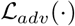 makes the recovered volume look realistic. The adversarial loss 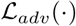 is computed from a trainable critic *D*(·), which works as a regularizer. We interpret this loss as a dynamic regularizer that is updated simultaneously with LISTA parameters during training. Note that only 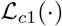 needs a labelled dataset, while 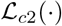 and 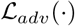 need LF images and unpaired 2P data, respectively.

The type of loss shown in Equation (7) has been first proposed as a supervised-learning technique for single image super-resolution [26]. Furthermore, learning regularizers to solve various standard inverse problems using stochastic gradient descent has been investigated in [27]. However, we highlight that our approach is a semi-supervised technique compared to [26]. Furthermore, the critic, or regularizer, is not pre-trained, as opposed to [27]. In our work, the adversarial regularizer is learned simultaneously with the generator (LISTA) by using the well-known adversarial training used for least squares GANs (LSGANs) [28], as depicted in Figure 7 (a). In the next section, we explicitly define the training loss and the architecture of the critic used for the experiments.

**Fig. 7.**
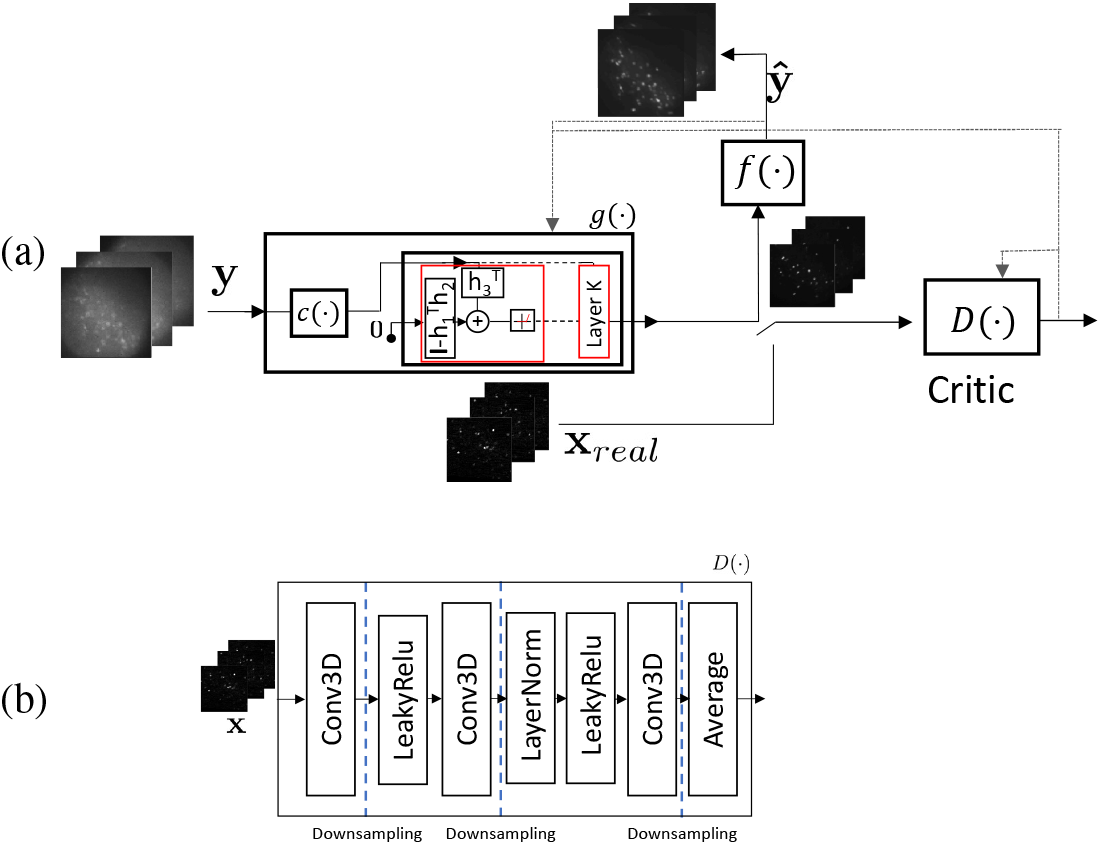
Training of our GAN architecture. In (a), we show how the LISTA network *g*(·) is trained using a content loss and an adversarial loss computed from a critic *D*(·). The content loss is computed using a few labelled data pairs, unlabelled LF data, and the known forward model *f*(·). In (b), we show the architecture of the critic *D*(·) designed following typical techniques for 3D GANs[25].

## VI. EXPERIMENTS AND RESULTS

In this section, we show the performance of our approach by imaging mouse brain tissue with both LF and 2P modalities (see supplementary material). We compare the performance of our method with state-of-the-art model-based and learningbased methods for reconstructing structural and functional LF data.

### A. Experimental Setup

The LF microscope is modelled as follows: numerical aperture = 1, refractive index = 1.33, wavelength = 514 *nm* for jGCaMP8f and 580 *nm* for TdTomato fluorophore, magnification = 25, microlens pitch =125 *μm*, microlens focal length = 1250 *μm*, tube lens focal length = 0.18 m, pixels per microlens =19 × 19.

We compare our method with two model-based approaches: ISRA, a variant of RL algorithm [15] and the ADMM approach proposed in [5]. Furthermore, we evaluate other learning-based approaches by adapting the LFMNet [10], HyLFM [11], and VCDNet [9] to work with our specifications based on the respective code made available online. To train our network and to evaluate model-based approaches, we use the conventional theoretical forward model used for reconstruction proposed by Broxton et al. [4].

We imaged mouse brain slices expressing TdTomato fluorescent protein with a 2P microscope to capture the spatial distribution of the network of neurons. The captured stack contains 80 planes taken at steps of 2 *μm*. Then, we generate 28 volumes with 53 slices each. The first volume includes planes 1 to 53. The second one contains planes 2 to 54, and so on. Similarly, we capture the corresponding LF stack with 28 images. Therefore, we have a training dataset with 28 training pairs taken from a single brain slice. For evaluation, we capture another dataset of the same size from a different brain sample.

In addition, we capture temporal LF sequences from samples labelled with genetically encoded calcium indicators (jGCaMP8f). We acquire 3 different temporal sequences with 500 LF images each. We took the first 80 LF images from each one for training. This additional training dataset only contains LF images without any 2P label. In our experiments, the 2P data is acquired only once and is not updated when evaluating different sample tissues.

Due to the dimensionality of the dataset, data augmentation is needed to alleviate data over-fitting. Specifically, we perform data augmentation by using random reflections on the *x, y*, and z-axis and axes swapping on the *x, y* dimension of the volume. We modify the LF data accordingly as well. Furthermore, we use patch-based training to reduce memory consumption.

### B. CNN Settings

To initialize the network, we pre-train it using only the labeled dataset. For this step, we use only the first content loss 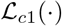 in Equation (7), which is chosen to be the normalized mean square error loss as follows:

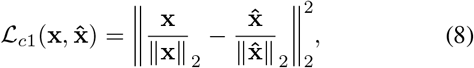

where **x** is the 2P 3D image and 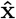 is the 3D image reconstructed by the network.

Once the network is initialized, all the losses in Equation (7), including the previous 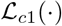 are considered in the minimization. The second content loss named 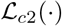 is given by the following equation:

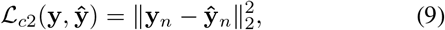

where the subscript *n* represents mean normalization, which is performed as follows:

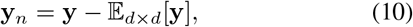

where the notation 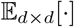 means that the expected value is computed for regions of size *d* × *d* pixels in the spatial dimensions. Specifically, each sub-aperture image of the LF **y** is divided into a grid of squares of size *d* × *d* pixels, then the expected value is subtracted from each square to obtain **y**_*n*_. This normalization allows focusing on reconstructing details in each sub-aperture image of the LF rather than background noise. In our experiments, the value d is experimentally chosen to be 8. For those familiar with wavelets, this procedure can be interpreted as subtracting the level 3 Haar approximation from each view in **y**. Thus, the loss only considers the horizontal, vertical, and diagonal details of levels 1,2, and 3.

The adversarial loss in Equation (7), 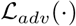 is determined by the discriminator *D*(·). The discriminator tries to assign different scores to the real 3D data and to the reconstruction from the network *g*(·). We name 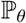 the probabilistic distribution of real 3D volumes and 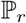 the distribution of 3D reconstructions from the network. Then, the adversarial loss is given by:

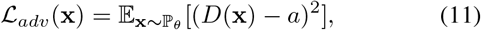

where *a* = 1 and the architecture of *D*(·) is depicted in Figure 7 (b). The discriminator was designed by following standard architectures used for 3D GANs [25]. The expected value 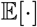 is approximated by computing the mean on a batch of volumes generated by LISTA. Finally, the discriminator is trained by using the loss

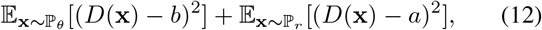

where *a* = 1, *b* = −1. As mentioned previously, the first expected value is computed on the batch of volumes generated with LISTA. Similarly, the second expected value is computed on batches of real data. Intuitively, the discriminator is trained to assign a 1 if the input has the 2P quality and a −1 if the input is generated by *g*(·) and does not have 2P quality. At the same time, *g*(·) tries to make *D*(·) to assign a 1 by improving the reconstruction quality. This training procedure is part of a standard technique proposed in [28] to train LSGANs.

### C. Reconstruction of Structural 3D images from Ligh Field images

In this section, we evaluate the performance of our method for reconstructing 3D images from a single LF image. We use unseen samples to measure the reconstruction performance by measuring the Peak Signal to Noise Ratio (PSNR) and Structural Similarity Index Measure (SSIM). The neurons in this sample are labelled with the TdTomato fluorescent fluorophore. Since this fluorophore is bright every time it is illuminated, these LF images only give structural information and do not show neuronal activity. As mentioned previously, we captured 28 LF images for training and 28 from a different sample for testing. Each LF image corresponds to a focal depth ranging from 0 to 54*μm*. From every LF image, we reconstruct a volume of size 321 × 321 × 53 voxels covering a range 533.3 × 533.3 × 104 *μm*^3^. The size of each LF image is 2033 × 2033 pixels.

The number of iterations used for model-based reconstruction must be chosen properly to avoid noise amplification. As mentioned in previous works [16], [7], [29], a typical empirical number of iterations used for ISRA is between 8 and 10. We fixed this value to 8 for both ISRA and ADMM. Our LISTA network comprises 6 unfolded iterations. Remarkably, 6 unfolded iterations of LISTA are enough to outperform competing methods.

The LISTA network achieves better average performance than other methods in a focal depth range of 54 *μm* in terms of PSNR and SSIM. In Table I, we show the performance for LF images taken at different focal depths; they are taken at steps of 2*μm*. The depth index in Table I increases as the depth increases. Note that all methods are affected by scattering as the depth increases. Even though all learning methods outperform model-based reconstruction approaches, our method achieves the best average performance in both PSNR and SSIM. The shown PSNR and SSIM are measured on the whole volume. In our experiments, the LFMNet achieved best performance among competing methods. Since the deep learning methods are trained with a very small dataset compared to the size of the dataset used in [10],[11],[9], their performance may be affected. In contrast, our method is more robust under this adverse condition. Furthermore, deep-learning methods are much faster than model-based approaches. Table II shows the average computational time to reconstruct a volume from a single LF image. All the methods were evaluated on a GeForce GTX 1080 Ti.

**TABLE I.**
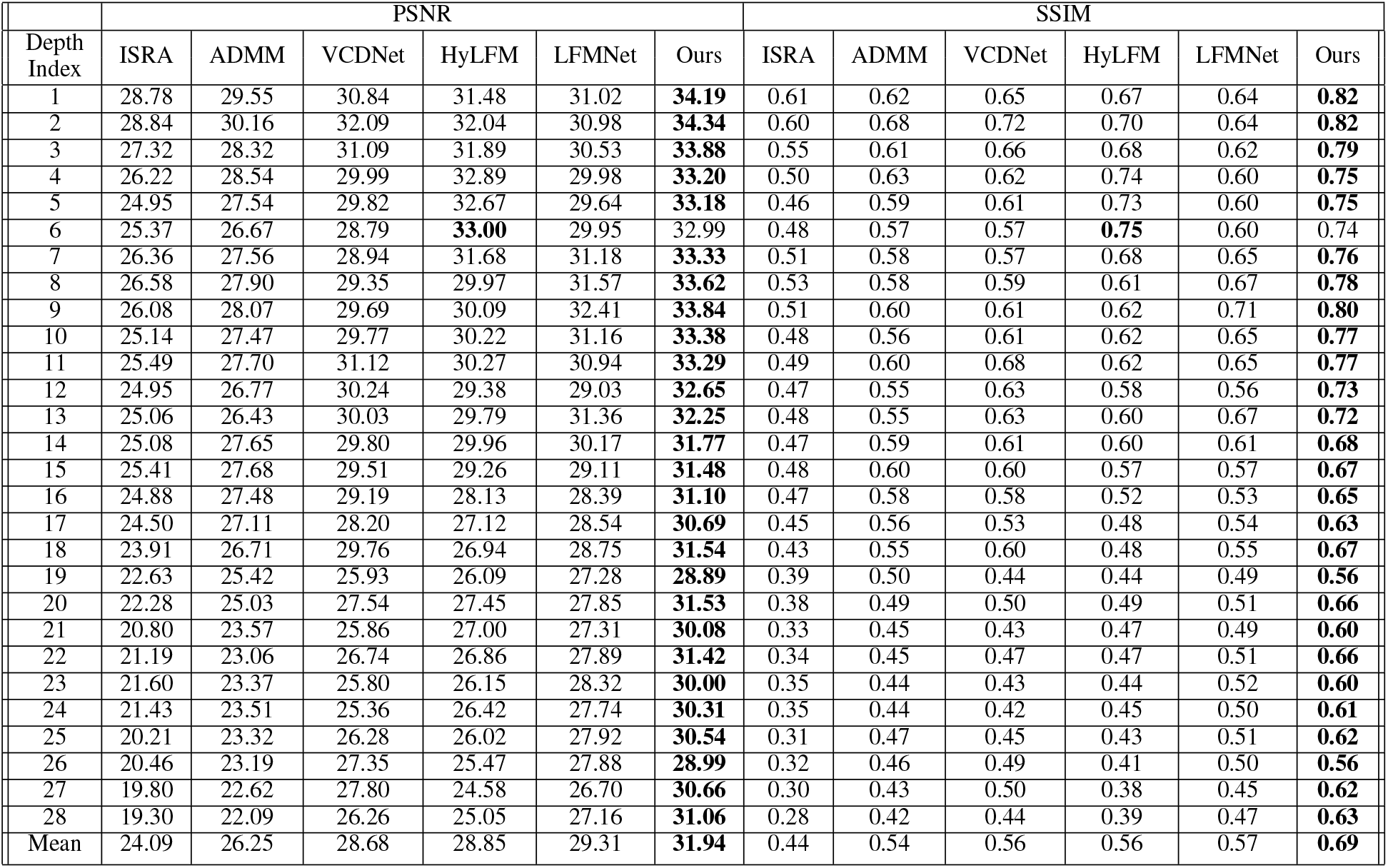
PSNR and SSIM for real lf data of neurons imaged using TdTomato fluorophore.

**Table 2.** Computational time.

|  | ISRA | ADMM | VCDNet | HyLFM | LFMNet | Ours |
| --- | --- | --- | --- | --- | --- | --- |
| Time (s) | 234.69 | 237.63 | 0.022 | 0.195 | 0.111 | 0.026 |

As shown in Figure 8, the LISTA network achieves better qualitative reconstruction performance than other methods. In Figure 8 (a) and (b), we show visual results for two different depths corresponding to index 1 and 28 in Table I, respectively. ISRA introduces square-like artifacts strongly present near the in-focus plane, approximately from *z* = −8 *μm* to *z* = 8 *μm* in part (a) and from *z* = 48 *μm* to *z* = 64 *μm* in (b). The ADMM can effectively remove these artifacts; however, both ISRA and ADMM are affected by background noise and scattering. As one goes deeper into the tissue, the modelbased methods are more affected by scattering. It is notable that learning methods achieve better performance than modelbased approaches and are less affected by noise. However, our approach is visually closer to the ground truth and achieves higher PSNR and SSIM than other learning methods. For instance, see plane *z* = 48 *μm*. Also, note in *z* = 64 *μm* that the LFMNet incorrectly reconstructs neurons from neighbour depths.

**Fig. 8.**
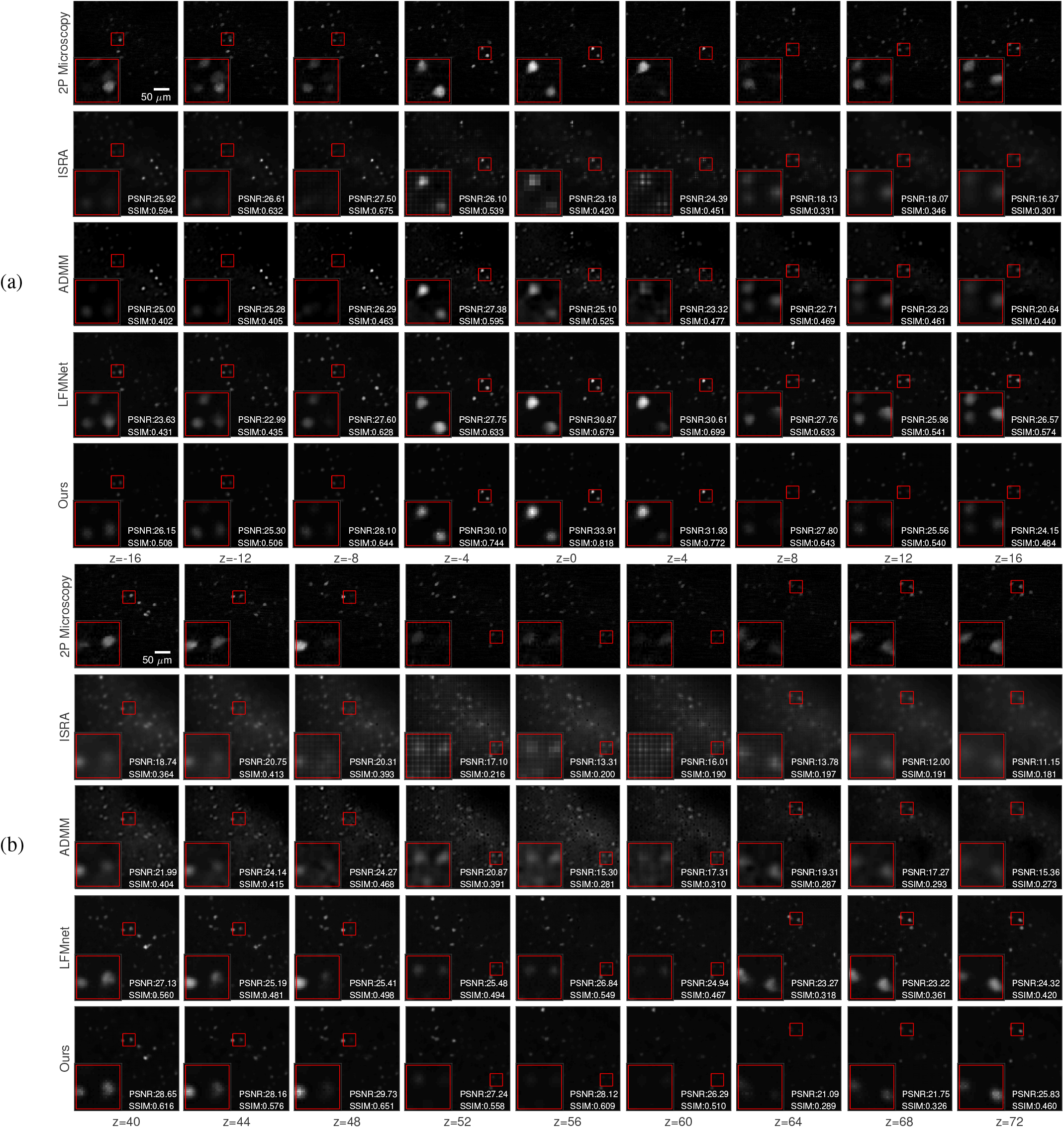
Reconstruction using real LF data from acute mouse brain slices expressing TdTomato fluorophore. In part (a), the first three rows show the 2P 3D image used as ground truth, and the reconstruction using two model-based approaches: ISRA and ADMM, respectively. Furthermore, in the next two rows we evaluate the state-of-the-art LFMNet proposed in [10] and we show our approach. We show several slices for different depths. This reconstruction corresponds to the performance shown in the first row in Table I. In part (b), we show performance for a LF image with a deeper focal depth, corresponding to the row 28 in Table I. The performance of all methods degrades when imaging deeper in the tissue. The shown PSNR and SSIM are measured for the whole plane at each depth. Measures on the whole volume are shown in Table I. All the distances are measured in *μm*. The settings used to capture both the LF image and 2P image are specified in Section VI.

### D. Reconstruction of volume time series from LF images

In this section, we evaluate the performance of our method for reconstructing a temporal sequence of 3D volumes from temporal sequences of LF images. The LF sequence captures the activity of neurons labelled with the jGCaMP8f calcium indicator at different times focused at a fixed focal depth. The jGCaMP8f is a fluorophore that increases its fluorescence intensity when the neurons fire. In this case, the ground truth data is unavailable since it is impossible to capture the activity of many neurons in 3-D with scanning-based techniques. We evaluate our approach on 3 LF sequences with 500 frames. As mentioned previously, only the first 80 LF images per sequence were used for training (with no labels), while the rest of LF images were unseen by the network.

Our LISTA network performs better than model-based approaches, while the state-of-the-art neural networks fail to reconstruct volumes for jGCaMP8f-labelled brain tissues. In Figure 9, we show the visual performance of ISRA, ADMM, and our method for reconstruction of one frame of the sequence (300th frame). We show two different samples in part (a) and (b). Even though the LFMNet [10] achieved satisfactory performance in the previous section, it generalizes poorly to the reconstruction of the temporal sequence due to the small training dataset. A more specific reason is that the use of a different fluorophore, the jGCaMP8f, implies samples with different noise levels and light sources with different wavelengths than those used for training. Figure 9 suggests that model-based methods are more robust under these adverse conditions than learning-based approaches, as mentioned in the introduction. However, model-based methods are heavily affected by scattering. In addition, ISRA introduces strong artifacts near the plane *z* = 0. In our approach, we exploit the knowledge of the forward model and the few available labels to achieve remarkable reconstruction performance.

**Fig. 9.**
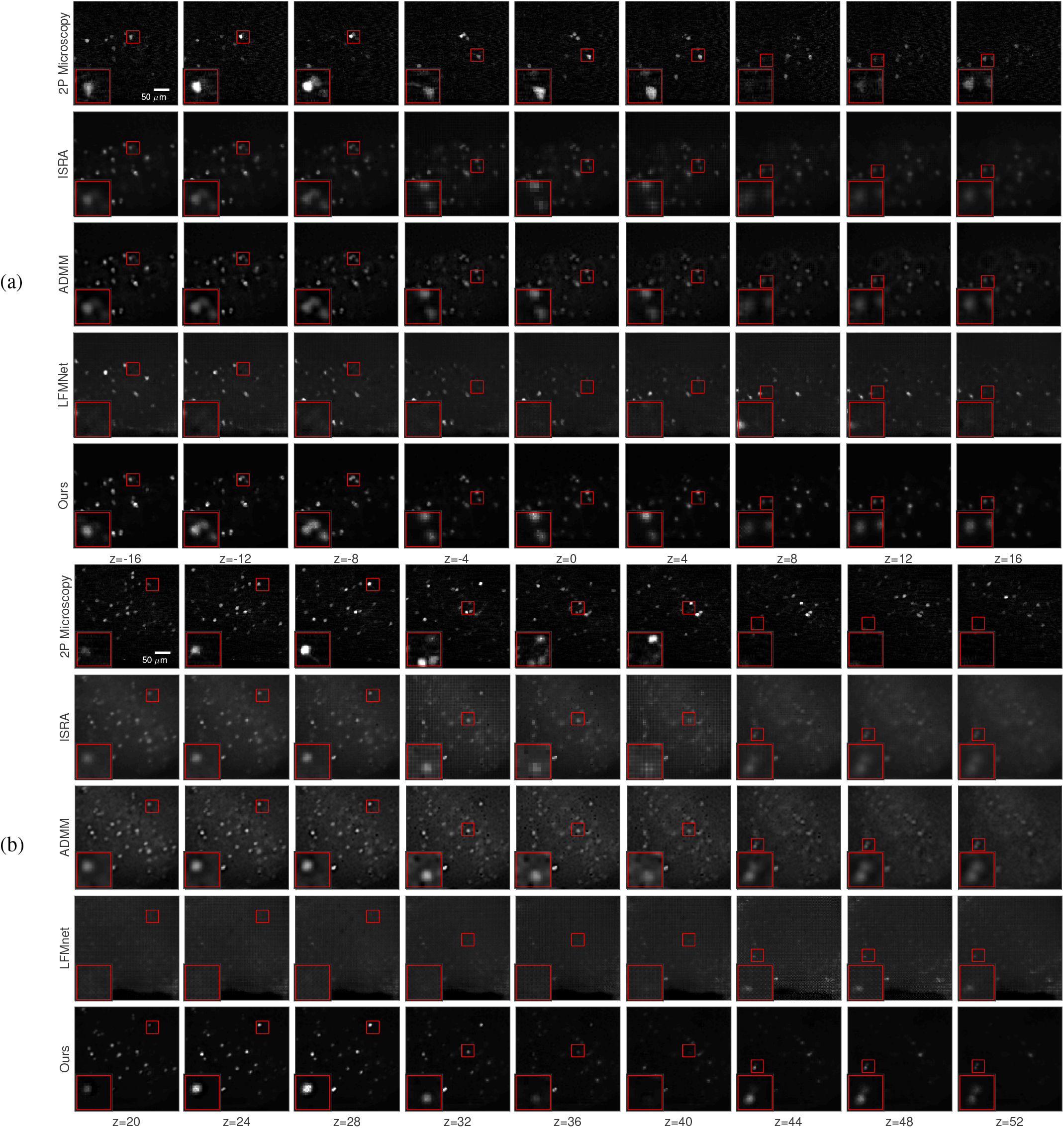
Reconstruction using real LF data from acute mouse brain slices expressing the calcium indicator jGCaMP8f. The indicator jGCaMP8f is suitable to perform functional imaging of neurons. The labels (a) and (b) indicate two different samples. The first row shows the 2P 3D image of the static tdTomato fluorophore used as a reference since the ground truth (2P jGCaMP8f volume) is unavailable. We show a particular reconstructed frame from a 500-frames sequence, see Section VI-D for details. We included the performance of the LFMNet [10] and two model-based approaches for comparison: ISRA and ADMM. We show different slices corresponding to different depths. All the distances are measured in *μm*. The settings used to capture both the LF image and the 2P 3D image are specified in Section VI.

Our training loss is designed to avoid amplifying noise from scattering. In Figure 10, we show the LF images synthesized from the reconstructed volumes. We display 3×3 views per LF image from the total 19 × 19 views. The re-synthesized LF image from ADMM shows that noise is reduced compared to the ISRA approach since the ADMM method imposes additional regularizers in the objective function. However, noise in the tissue region is still significant. Our approach greatly reduces noise while reconstructing every neuron footprint in the ground truth LF image. We achieve this remarkable performance due to the adversarial regularizer and the specialized content losses used for training.

**Fig. 10.**
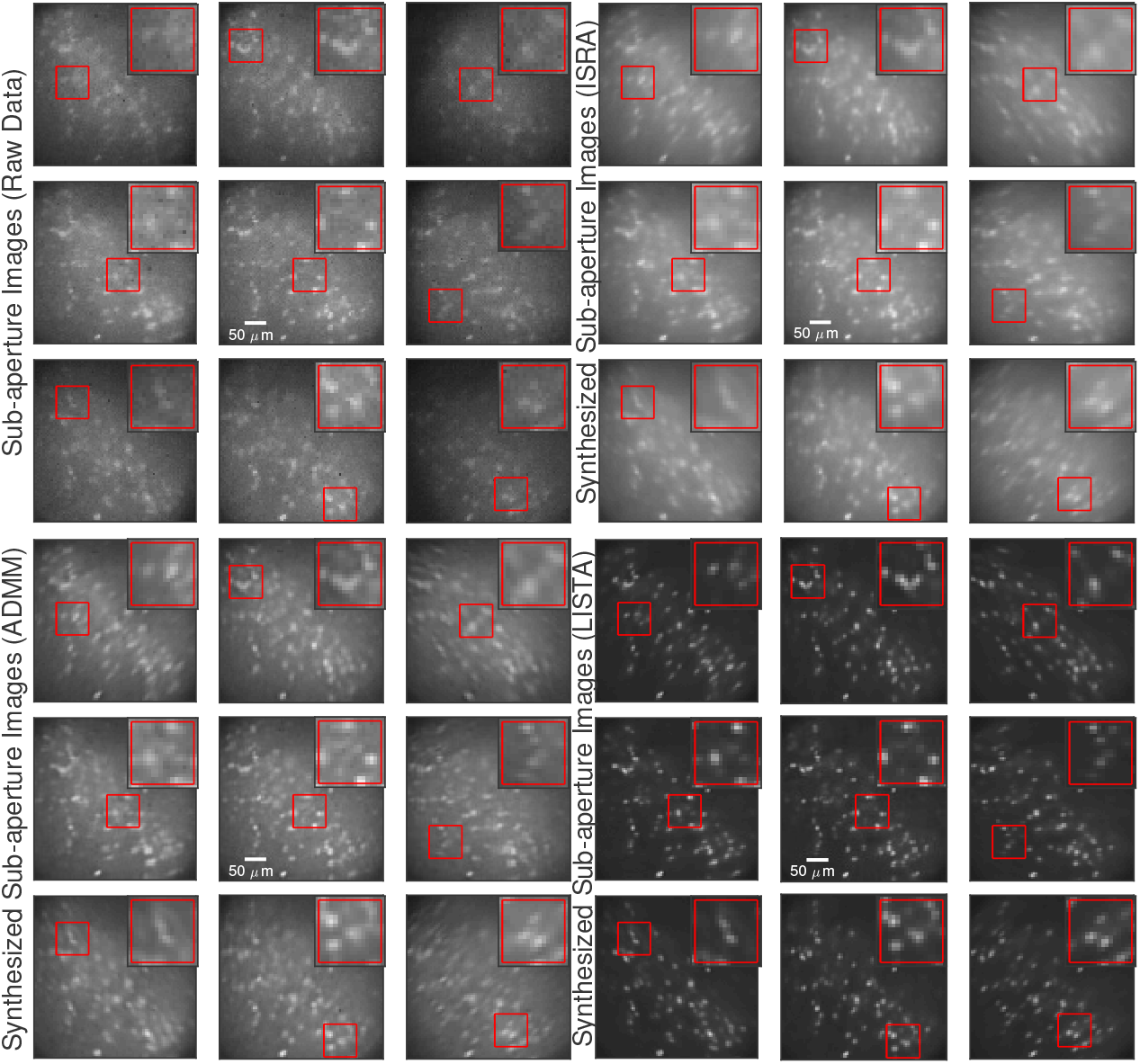
Visual comparison of sub-aperture images. We show 9 different sub-aperture images (views) for 4 different cases: views from the raw LF data, LF synthesised from ISRA reconstruction, LF synthesised from ADMM method, and LF synthesised from our LISTA approach. Both model-based methods reconstruct noisy regions in the sub-aperture images that do not carry any meaningful information, which translates into a noisy 3D reconstruction. On the other hand, our approach can accurately reconstruct every neuron footprint while significantly reducing noise from scattering.

Our approach provides a new powerful tool to study fast temporal evolution of neurons in mammalian brain tissue. In Figure 11, we show reconstruction of temporal evolution of neurons. We show three brain samples in parts (a), (b), and (c). For this experiment, we perform reconstruction from LF sequences of 500 frames. Therefore, we reconstruct 500 3D volumes per LF sequence with the same size as in previous experiments. Since the imaging rate is 50 Hz, we show 10 seconds of neuronal activity. Note how neurons located at different positions in the 3D space show different activity.

**Fig. 11.**
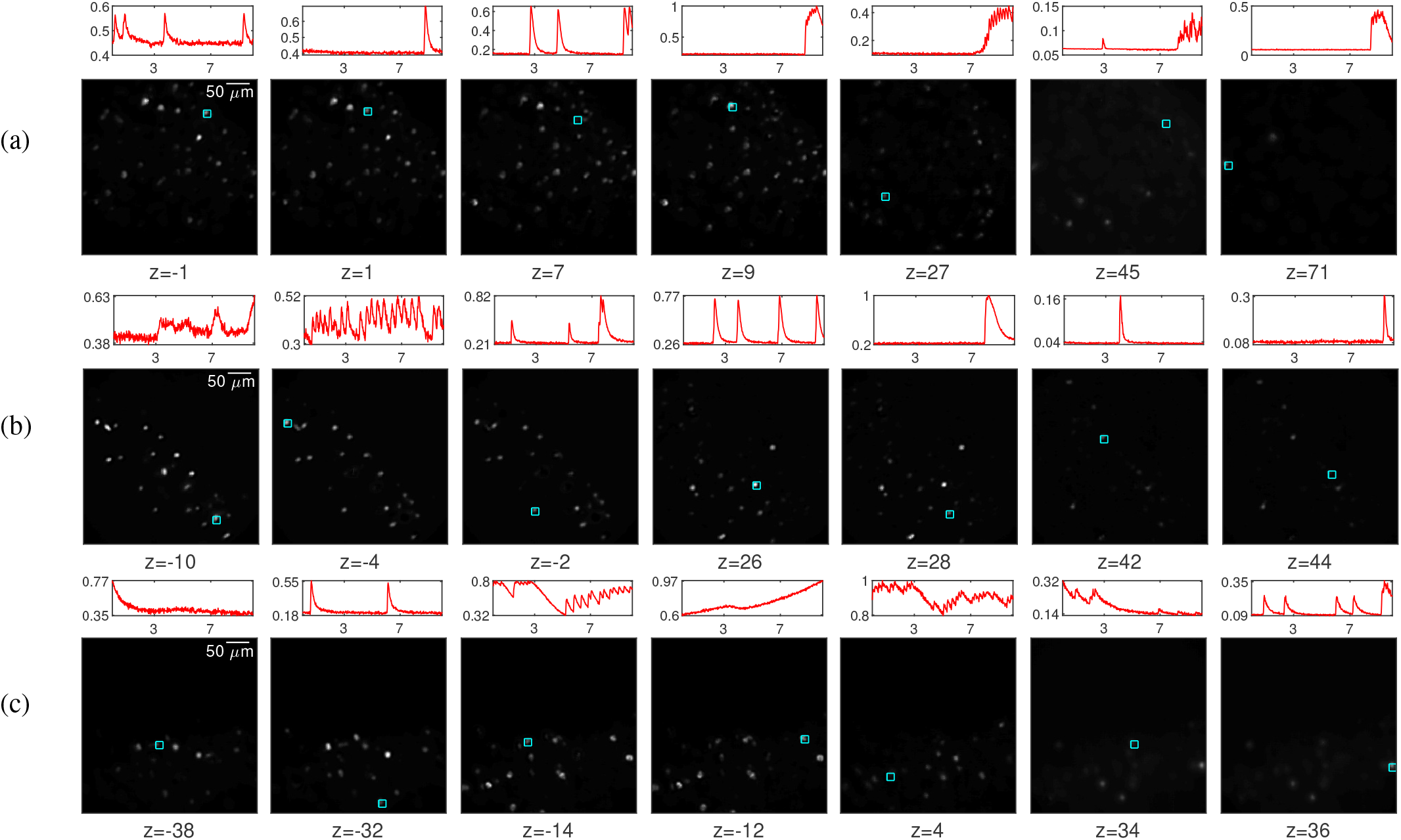
Reconstruction of the temporal evolution of neuron activity in brain tissue. Our LISTA network allows fast reconstruction of volume time series from LF data of mammalian brain tissue expressing the calcium indicator jGCaMP8f. The labels (a), (b), and (c) indicate three different sample tissues. We show several slices containing active neurons, the neuron of interest is marked in cyan. On top of each slice, we show the temporal evolution of the neuron. The horizontal axis shows time in seconds, the vertical axis shows the normalized intensity. The normalization is performed per sample between the selected neurons. Our method reconstructs 500 frames (500 3D images) in this experiment, a computationally demanding task with conventional model-based methods. The LF imaging rate is 50 Hz. All the distances are measured in μm. Other additional settings are specified in Section VI.

## VII. Conclusion

We have introduced a physics-driven deep neural network to reconstruct 3D volumes from LF sequences. The architecture of the network is based on the observation that labelled neurons in tissues are sparse, which naturally leads to an architecture based on unfolding the ISTA algorithm. Moreover, we show how the forward model of a LF microscope can be described using CNNs, and this physics-driven architecture is also included in the network. Finally, we use GANs and exploit the theoretical forward model to train our network with a small labelled dataset.

We show that our method achieves better reconstruction quality for reconstructing temporal LF sequences imaged with the jGCaMP8f indicator than other state-of-the-art methods. None of the competing learning-based methods can perform this task. Furthermore, we also showed better performance than standard model-based and learning-based methods in terms of PSNR and SSIM for reconstructing structural 2P volume imaged using the TdTomato fluorophore. Although LFM cannot penetrate as deeeply into scattering tissue as functional 2P imaging, our 2P-enhanced LFM strategy enables light-efficient volume acquisition at fast rates, essential to capturing neuronal dynamics transduced by fast sensors such as jGCaMP8f [30].

Our work offers a practical method to perform 3D reconstruction for LF microscopy under adverse acquisition conditions when imaging mammalian brain tissue. We believe that the proposed method could be helpful even beyond this specific scenario, and it could inspire similar solutions for other types of inverse problems.

## Supporting information

Experimental setup

## Acknowledgment

The authors would like to thank Dr. Yu Liu for preparing the plasmids and for assisting with the fluorophore transfection.

## Notes

This work is in part supported by BBSRC BB/R009007/1 and in part by the Royal Academy of Engineering under the RAEng Research Fellowship scheme (RF1415/14/26). Flavie Lesept is supported by Marie Skłodowska-Curie Fellowship (707478). Herman Verinaz-Jadan was in part supported by SENESCYT.

### Competing Interest Statement

The authors have declared no competing interest.

