## Supplementary material for "Physics-based Deep Learning for Imaging Neuronal Activity via Two-photon and Light Field Microscopy": Experimental setup

### I. Supplemental Material

#### A. Experimental Setup

We performed multimodal light field and two-photon imaging of mouse live brain slices in which layer $2/3$ excitatory neurons were co-transfected with the fluorophores jGCaMP8f (calcium indicator) and tdTomato (static, morphology label). Imaging was performed with a custom-built microscope featuring coaligned light-field and two-photon imaging paths. Specifically, imaging was performed with a custom-built epifluorescence microscope with an MLA (125 $\mu m$ pitch, ff/10, RPC Photonics) placed at the imaging plane of a $25\times$, numerical aperture (NA) =1.0 water immersion objective lens (XLPLN25XSVMP, Olympus), and 180-$mm$ tube lens (TTL180-A, Thorlabs). The MLA was imaged onto a scientific complementary metal–oxide–semiconductor (sCMOS) camera (ORCA Flash 4 V2 with Camera Link, 2048 $\times$ 2048 pixels, 6.5 $\mu m$ pixel size, Hamamatsu) with a 1:1 relay macro lens (Nikon 60 $mm$ f2.8 D AF Micro Nikkor Lens). The MLA was aligned following the advice in [1]. TdTomato fluorescence was excited with a 530 $nm$ LED, and jGCaMP8f fluorescence was excited by a 470 $nm$ LED, both powered by an OptoLED current driver (P1110/002/000, Cairn Research). Light-field images were collected at $50 - 100$ frames per second using Micromanager 2.0-gamma [1].

*1) Light-field modality specifications:* Imaging was performed with a custom-built epifluorescence microscope with an MLA ($125\mu m$ pitch, ff/10, RPC Photonics) placed at the imaging plane of a $25\times$, numerical aperture (NA) =1.0 water immersion objective lens (XLPLN25XSVMP, Olympus), and 180-$mm$ tube lens (TTL180-A, Thorlabs). The MLA was imaged onto a scientific complementary metal–oxide–semiconductor (sCMOS) camera (ORCA Flash 4 V2 with Camera Link, $2048 \times 2048$ pixels, 6.5 $\mu m$ pixel size, Hamamatsu) with a 1:1 relay macro lens (Nikon 60 $mm$ f2.8 D AF Micro Nikkor Lens). The MLA was aligned following the advice in [1]. TdTomato fluorescence was excited with a 530 $nm$ LED, and jGCaMP8f fluorescence was excited by a 470 $nm$ LED, both powered by an OptoLED current driver (P1110/002/000, Cairn Research). Light-field images were collected at $50 - 100$ frames per second using Micromanager 2.0-gamma [1].

*2) Two-photon modality specifications:* The two-photon imaging laser (Coherence Monaco 1035-40-40, central wavelength 1035 $nm$, pulse frequency 10 MHz) was introduced between the light-field imaging system objective and tube lens with a dichroic mirror (DI03-R785-T3 $25 \times 36 \times 3$ $mm$, Semrock). The laser was focused by a $f$=30 $mm$ scan lens, $f$=300 $mm$ tube lens, and the common objective lens (XLPLN25XSVMP, Olympus). The laser was scanned laterally (x,y) by two 3 $mm$ mirrors (6M2003S-S, Cambridge Technologies) driven by high power servo electronics (671315K-1HP, Cambridge Technologies). The plane of focus was adjusted by moving the objective with a stepper motor (SliceScope, Scientifica). tdTomato fluorescence was collected by a $50 \times 70 \times 2$ $mm$ dichroic mirror (T750lpxrxt-UF2, Chroma) positioned directly above the objective back aperture only during two-photon imaging. The objective back pupil was conjugated and demagnified onto the active area of a photomultiplier tube with integrated transimpedance amplifier (PMT, Hamamatsu H10722-20-10MHz). The tdTomato fluorescence passed through a 750 $nm$ shortpass filter (Semrock FF01-750/SP-25) before the PMT. A National Instruments PCI-6110 drove the scan mirrors and digitized the PMT/amplifier output through ScanImage version 3.8 software.

*3) Fluorophore transfection:* Mouse layer 2/3 cortical neurons were transfected via in-utero electroporation (IUE) with soma-targetedw [2] jGCaMP8f [3] (pAAV-CAG-RiboGCaMP8f) and tdTomato (pCAG-tdTomato [4], Addgene). On embryonic day (E)15.5 timed-pregnant female CD-1 mice (Charles River UK) mice were deeply anaesthetized with 2% isoflurane. Uterine horns were exposed and periodically rinsed with warm sterile PBS. Plasmid DNA, 1–2 $\mu g$ total at a final concentration of 1 $\mu g/\mu l$ (a 6:1 ratio of jGCaMP8f:tdTomato) diluted in sterile PBS was injected into the lateral ventricle of one cerebral hemisphere of an embryo. Five voltage pulses ($50V$, $50ms$ duration, $1Hz$) were delivered using 5-mm round plate electrodes (ECM 830 electroporator, Harvard Apparatus), with the anode or cathode placed on top of the skull to target the cortex or hippocampus, respectively. Electroporated embryos were placed back into the dam, and allowed to mature to delivery. Brain slices were prepared from electroporated mice at postnatal day (P)12–P30.

*4) Brain slice preparation:* This study was carried out in accordance with the recommendations of the UK Animals (Scientific Procedures) Act 1986 under Home Office Project and Personal Licenses (project license 70/9095). 400-$\mu m$ slices were prepared from $12 - 30$ day old mice Slices were cut in choline chloride cutting solution containing (in $mM$): 110 choline-Cl, 25 $NaHCO_3$, 20 glucose, 2.5 KCl, 1.25 $NaH_2PO_4$, 0.5 $CaCl_2$ and 7 $MgSO_4$. After cutting, the slices were transferred to a solution containing (in $mM$) 125 NaCl, 25 $NaHCO_3$, 20 glucose, 2.5 KCl, 1.25 $NaH_2PO_4$, 2 $MgSO_4$, 2 $CaCl_2$, adjusted 300 to 310 mOsm/kg, pH 7.3 to 7.4 with HCl at $36°C$. All solutions were oxygenated with $95\%O_2/5\%CO_2$.
